# Tripartite AAV Systems for *EYS* Retinal Gene Therapy

**DOI:** 10.64898/2025.12.03.692187

**Authors:** Kun-Do Rhee, Poppy Datta, Clairissa Baccam, Seongjin Seo

## Abstract

Mutations in the *Eyes Shut Homolog* (*EYS*) gene are a leading cause of autosomal recessive retinitis pigmentosa, a progressive retinal degenerative disease for which no effective treatment currently exists. However, the large size of the *EYS* coding sequence (∼9.4 kb) exceeds the packaging limit of adeno-associated virus (AAV) vectors, posing a major barrier to gene replacement therapy. To address this challenge, we developed a tripartite AAV vector system that enables delivery and reconstitution of the full-length *EYS* gene using a Cre-lox-based unidirectional DNA recombination strategy, Uni-STAR (Uni-directional and Site-specific Transgene Assembly by Recombination). The system consists of three AAV constructs carrying discrete *EYS* segments flanked by engineered, non-compatible lox sites that drive ordered and unidirectional recombination in target cells. We validated this system *in vitro* by demonstrating successful reconstitution and expression of full-length EYS protein in HEK293T cells. *In vivo*, subretinal co-injection of the three AAV vectors into mouse eyes led to precise reconstitution and expression of full-length EYS protein in the retina. These findings establish the feasibility of using a tripartite AAV system to deliver the complete *EYS* gene and provide a foundation for future therapeutic development targeting *EYS*-associated retinal degenerations.

## Introduction

Mutations in the *Eyes Shut Homolog* (*EYS*) gene are a major cause of autosomal recessive retinitis pigmentosa (RP), accounting for 5-10% of all cases.^1–7^ Clinically, EYS-associated RP typically manifests with night blindness in the second or third decade of life. This is followed by progressive peripheral visual field constriction and eventual loss of central vision. However, phenotypic heterogeneity exists: some patients exhibit early macular involvement or cone degeneration, and autosomal recessive cone-rod dystrophy (CRD) is also associated with *EYS* mutations.^8–10^ Although no therapy currently exists, the relatively slow progression of photoreceptor degeneration offers a window of opportunity for therapeutic intervention prior to irreversible retinal damage.

The human *EYS* gene, an ortholog of *Drosophila melanogaster eyes shut* (also known as *spacemaker* (*spam*))^11,12^, is located on chromosome 6p12. It spans more than 2 Mb and includes 44 exons. Four isoforms are expressed in the human retina, with isoforms 4 and 1 encoding the longest proteins (9,498 and 9,435 bp, respectively).^13^ Both isoforms contain 20 epidermal growth factor (EGF)-like domains in the N-terminal half and five Laminin G-like domains interspersed with seven additional EGF-like motifs in the C-terminal region.

In *Drosophila*, Eys is required for the formation of the matrix-filled inter-rhabdomeral space.^11,12^ While rodents lack *EYS* orthologs, zebrafish express *eys* in photoreceptor cells, with protein localization near the connecting cilium.^14,15^ Eys deficiency in zebrafish causes mislocalization of key outer segment proteins and subsequent photoreceptor degeneration.^14–16^ More recently, retinal organoids were generated from patient-derived induced pluripotent stem cells (iPSCs), and G protein-coupled receptor kinase 7 (GRK7) was shown to be mislocalized in *EYS* mutant retinal organoids.^17^ However, the precise molecular function of EYS and the pathomechanisms of retinal degeneration by EYS mutations remain poorly understood.

Gene replacement therapy using adeno-associated virus (AAV) vectors is a promising option for treating inherited retinal diseases. AAV is currently the preferred gene transfer vector for retinal gene therapy, owing to its well-established safety, long-term transgene expression, and remarkable efficiency in transducing post-mitotic retinal cells, including photoreceptors. However, the most significant limitation of AAV vectors is their restricted packaging capacity, which is constrained to approximately 4.7 kb of single-stranded DNA. This size constraint precludes the delivery of many large genes, including *EYS*, using a conventional single AAV vector.

We have recently reported a new strategy, termed Uni-STAR (<u>Uni</u>-directional and <u>S</u>ite-specific <u>T</u>ransgene <u>A</u>ssembly by <u>R</u>ecombination), to deliver large genes using up to four AAV vectors.^18^ In Uni-STAR, target genes are split into two to four segments, each carried by a separate AAV vector, and reconstituted via a unidirectional, site-specific DNA recombination system consisting of engineered lox sites and Cre recombinase. In the present study, we applied this strategy to *EYS*, developing a tripartite AAV vector system capable of delivering and reconstituting the full-length *EYS* gene. We demonstrate successful EYS protein expression both *in vitro* and *in vivo* in the mouse retina, laying the foundation for therapeutic applications.

## Materials and Methods

### Plasmid Construction

A full-length human *EYS* cDNA clone (pcDNA3.1-hEYS; NM_001142800.2) was obtained from GenScript and used as a template for PCR amplification. The coding sequence (9,435 bp) was divided into three segments (1,644 bp, 3,903 bp, and 3,888 bp), PCR-amplified using Q5 High-Fidelity DNA polymerase (New England Biolabs), and cloned into AAV shuttle plasmids (pAAV-GRK1p-15-inCREv1 (with T2A), pAAV-GRK1p-15-IRES-inCRE (with IRES), pFBAAV-17-mid-1522, and pFBAAV2-1722-pA)^18^ using a GenBuilder Cloning kit (GenScript). The resulting constructs were pAAV-GRK1p-EYS-E1-15-inCRE, pAAV-GRK1p-EYS-E1-15-IRES-inCRE, pFBAAV-17-EYS-E2-1522, and pFBAAV2-1722-EYS-E3-pA, respectively. Primer sequences are available upon request.

### Cell Culture and Transfection

HEK293T/17 cells (ATCC #CRL-11268) were cultured and transfected as previously described.^18^

### Animals and Subretinal Injection

Wild-type C57BL/6J mice (strain #: 000664) were acquired from the Jackson laboratory. All procedures were approved by the Institutional Animal Care and Use Committee (IACUC) of the University of Iowa and conducted in accordance with the recommendations outlined in the Guide for the Care and Use of Laboratory Animals of the National Institutes of Health. Mice were kept on a 12-hour light/dark cycle with *ad libitum* access to standard mouse chow.

For subretinal injection, AAV-*EYS* vectors (serotype: AAV8) were produced by the University of Iowa Viral Vector Core. On the day of injection, AAV vectors were thawed on ice and mixed 1:1:1 (3x10^9^ genome copies (GC)/μl per vector). Both male and female mice at 1 month of age were anesthetized with a ketamine/xylazine mixture (87.5 mg/kg ketamine, 12.5 mg/kg xylazine), and 10% povidone-iodine and 1% tropicamide solutions were applied. Under a Zeiss SteREO Discovery.V8 dissecting microscope, eyeballs were slightly pulled out using forceps, and a small hole was made using 30-gauge needle near limbus. Limbal-approach trans-retinal subretinal injections were performed using a NanoFil syringe attached to a blunt-end 35-gauge needle (World Precision Instrument) as previously described,^19^ delivering 1 μl of the vector solution. After injections, an antibiotic/steroid ophthalmic ointment (neomycin and dexamethasone) was applied. Antisedan (Zoetis) was administered right after injection to facilitate recovery. Animals were excluded from follow-up analysis if no obvious blebs were observed or if significant hemorrhage occurred at the time of injection.

### Viral DNA Extraction from Mouse Retinas and PCR

Mice were euthanized by CO_2_ asphyxiation followed by cervical dislocation, and eyes were enucleated. Anterior segments were removed with micro-dissecting scissors, and posterior segments were homogenized in 1 ml of TRIzol Reagent (Invitrogen) using a Polytron PT 1200E homogenizer (Kinematica). Episomal viral DNA was extracted following the manufacturer’s RNA extraction protocol and subsequently used as a template for PCR with GoTaq G2 Flexi DNA polymerase (Promega). The following junction-specific primers were used to detect recombination between vector segments: for the 5’-middle junction, F1 (5’-CCAAGATAAAGGTCCTGCTCAA-3’) and R1 (5’-CAGAGGCCATGCACTGATATAC-3’); and for the middle-3’ junction, F2 (5’-CGTCTTCCTCCATGTCTGTAAT-3’) and R2 (5’-CAGAAGTCCATAGGAGCTGAAG-3’).

### Protein extraction, SDS-PAGE, and immunoblotting

Proteins were extracted from HEK293T/17 cells and mouse retinas as previously described.^18^ Extracted proteins were loaded on 3-8% NuPAGE Tris-Acetate gels (Invitrogen), transferred onto nitrocellulose membranes (BioRad), and immunoblotted following standard protocols. Antibodies used are rabbit anti-EYS (Invitrogen, PA5-55507), rabbit anti-Cre (Cell Signaling, 5036), mouse anti-β-actin (Sigma, A1978), horse radish peroxidase (HRP)-conjugated anti-mouse IgG (Cell Signaling, 7076), and HRP-conjugated anti-rabbit IgG (Cell Signaling, 7074). Detection was performed using SuperSignal West Dura Extended Duration Substrate (Thermo Scientific) and imaged with a ChemiDoc Imaging system (Bio-Rad).

## Results

### Design of a Tripartite AAV Vector System for Full-Length EYS Gene Delivery

To enable AAV-mediated delivery of the full-length *EYS* coding sequence (CDS; 9,435 bp, NM_001142800.2, isoform 1), we engineered a tripartite vector system comprising three AAV constructs, each carrying a distinct segment of the *EYS* CDS (**Figure 1A**). These segments were designed to be seamlessly reconstituted in target cells via Cre-mediated, unidirectional DNA recombination.

**Figure 1.**
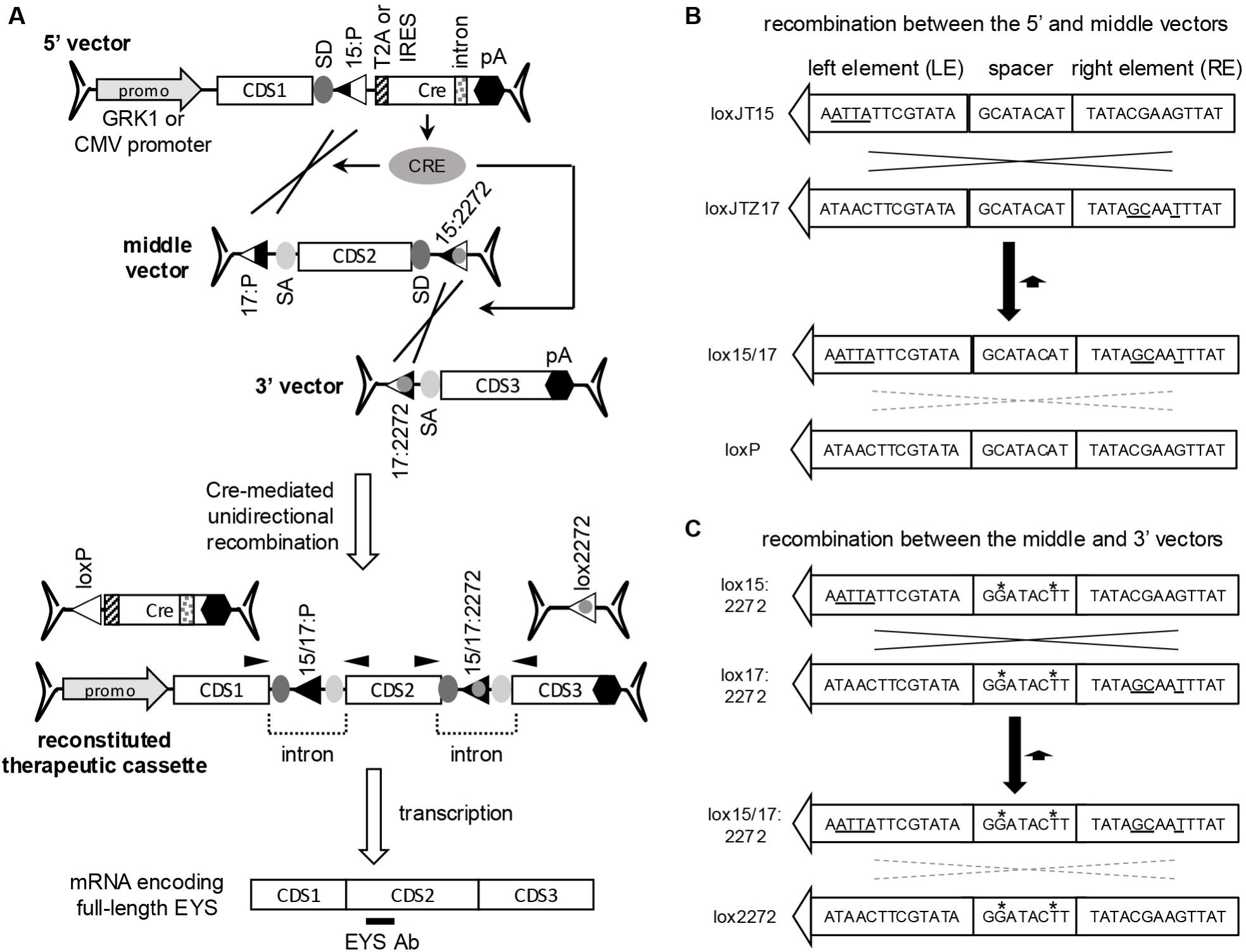
Strategy for reconstituting the full-length *EYS* gene using a triple AAV vector system. (A) Overview of the Cre-mediated recombination system. The *EYS* coding sequence (CDS) was split into three segments (CDS1, CDS2, and CDS3) and incorporated into three separate AAV vectors: 5’, middle, and 3’. In the reconstituted cassette, splice donor (SD) and splice acceptor (SA) sites flanking the CDS fragments form functional introns. Black arrowheads (F1, R1, F2, and R2) on the reconstituted therapeutic cassette denote the binding sites of PCR primers used to amplify the reconstituted junctional regions. The location of the EYS antibody immunogen (residues C795-R861) is indicated by a black bar. Lox site variants were labeled as follows: 15:P (loxJT15 with a loxP spacer), 15:2272 (loxJT15 with a lox2272 spacer), 17:P (loxJTZ17 with a loxP spacer), 17:2272 (loxJTZ17 with a lox2272 spacer), 15/17:P (loxJT15/JTZ17 LE/RE double mutant with a loxP spacer), 15/17:2272 (loxJT15/JTZ17 LE/RE double mutant with a lox2272 spacer). Additional abbreviations: IRES, internal ribosomal entry site; pA, polyadenylation signal; T2A, T2A “self-cleaving” peptide. (B) Lox sites for recombination between the 5’ and middle vectors. This recombination is mediated by loxJT15 and loxJTZ17 sites, which are loxP variants containing mutations (underlined) in either the left element (LE) or the right element (RE). Cre-mediated recombination between these two sites produces one LE/RE double mutant (lox15/17) and one canonical loxP site. The double mutant lox15/17 site binds Cre with very low affinity, preventing the reverse reaction. (C) Lox sites for recombination between the middle and 3’ vectors. Lox15:2272 and lox17:2272 are LE and RE mutant lox sites, respectively, with a lox2272 spacer (GGATACTT) instead of a loxP spacer (GCATACAT). Mutations in the lox2272 spacer are marked by asterisks (*).

The 5′ vector contains a photoreceptor-specific GRK1 promoter (or a ubiquitously active CMV promoter for *in vitro* validation), followed by the first *EYS* segment (1,644 bp), a splice donor (SD) site, a modified lox site (loxJT15), and a bicistronic Cre recombinase expression cassette (T2A “self-cleaving” peptide or IRES-based) with an embedded intron to prevent prokaryotic expression, and a bovine growth hormone (BGH) polyadenylation (polyA) signal. The middle vector carries a loxJTZ17 site (compatible with loxJT15 in the 5’ vector), a splice acceptor (SA), the second *EYS* segment (3,903 bp), a second SD, and a lox15:2272 hybrid lox site (compatible with the lox17:2272 site in the 3’ vector). The 3′ vector contains a lox17:2272 hybrid site, a SA, the third *EYS* segment (3,885 bp), and a BGH polyA signal.

The loxJT15 and loxJTZ17 pair contains the canonical loxP spacer (GCATACAT) (**Figure 1B**), whereas the lox15:2272 (hybrid of loxJT15 and lox2272) and lox17:2272 (hybrid of loxJTZ17 and lox2272) pair has the lox2272 spacer (GGATACTT) (**Figure 1C**).^18,20,21^ Because these two spacers are non-compatible, cross-recombination between the two pairs does not occur. This incompatibility prevents excision of the floxed *EYS* second segment from the middle vector as well as from the reconstituted expression cassette in the presence of Cre. Additionally, loxJT15 and loxJTZ17 are reaction equilibrium-modifying variants of loxP.^20^ These lox sites have mutations in one of the two 13-bp inverted repeats (left element (LE) and right element (RE)), which serve as Cre binding sites. The use of these modified lox sites suppresses reverse reactions, rendering the recombination events essentially irreversible.

Since lox sites are directional (i.e., asymmetric) and Cre-mediated recombination is sequence-specific, this design ensures the ordered and orientation-correct assembly of the three *EYS* segments. The SD and SA sites in the reconstituted expression cassette facilitate RNA splicing of the transcribed pre-mRNA. This splicing process removes any intervening non-coding sequences (including the recombined lox sites), thereby generating a single, continuous mRNA molecule that encodes the full-length EYS protein. Furthermore, the use of a bicistronic element obviates the need for a separate promoter for Cre expression, and the placement of a lox site before the bicistronic element allows self-inactivation of Cre after the reconstitution of therapeutic cassettes by separating the Cre CDS from its promoter.

### Reconstitution and Expression of Full-Length EYS in 293T Cells

To validate the feasibility of the tripartite vector system *in vitro*, HEK293T cells were transfected with the three AAV constructs (**Figure 2**). The 5′ vector employed the CMV promoter and T2A peptide for bicistronic Cre expression. Control conditions included individual plasmids (lanes 1-3) and a full-length *EYS* expression construct (lane 5). Western blot analysis using an anti-EYS antibody raised against residues C795-R861 revealed a ∼350 kDa band corresponding to full-length EYS only in cells transfected with all three vectors simultaneously (lane 4). The reconstituted EYS band co-migrated with the band observed in cells transfected with the full-length *EYS* expression plasmid, confirming accurate assembly and translation. Cre protein was detected in lysates from cells receiving the 5′ vector. Expression of β-actin was consistent across all lanes, serving as a loading control. These results provide strong proof-of-principle that the tripartite AAV system can reconstitute the full-length *EYS* CDS and produce correctly sized protein in mammalian cells.

**Figure 2.**
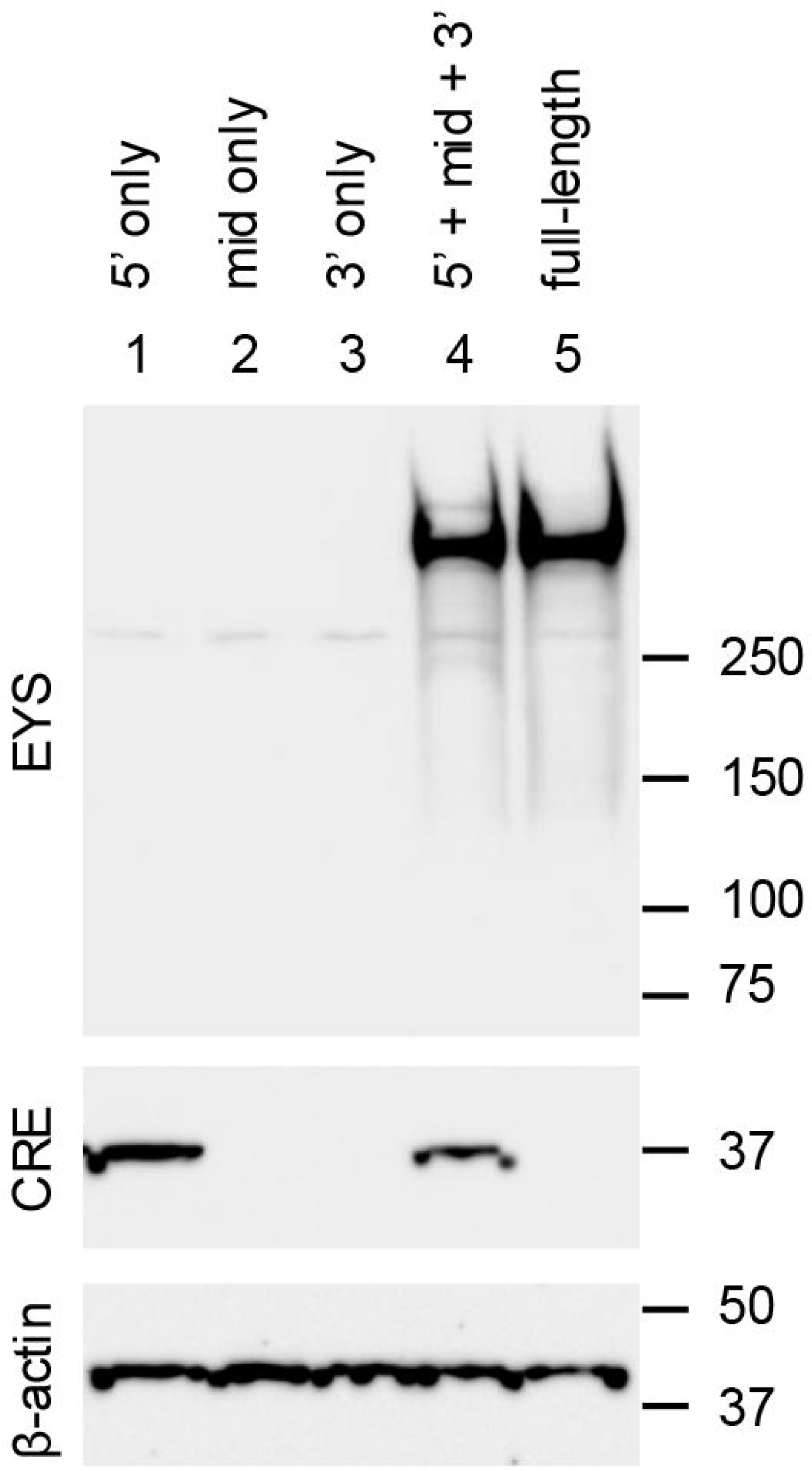
*In vitro* reconstitution and expression of full-length EYS protein in HEK293T cells. HEK293T/17 cells were transfected with plasmids encoding individual (lanes 1-3) or all three (lane 4) segments of the tripartite AAV-*EYS* system. A full-length *EYS* expression plasmid served as a positive control (lane 5). Lysates were collected three days post-transfection and subjected to SDS-PAGE followed by Western blot analysis using an anti-EYS antibody. β-actin served as a loading control. Molecular weight markers are indicated on the right.

### Reconstitution and Expression of Full-Length EYS in Mouse Retinas

To evaluate the feasibility of the tripartite vector system *in vivo*, we used mice as a cost-effective surrogate model despite the absence of an *EYS* ortholog in their genome. The tripartite AAV-*EYS* vectors were packaged into AAV8 capsids and delivered into the subretinal space of 1-month-old wild-type C57BL/6J mice. Two versions of the 5′ vector were tested—one containing a T2A-based Cre cassette and another containing an IRES-based cassette—both driven by the GRK1 promoter for photoreceptor-specific expression. Each vector was injected at 3 × 10 genome copies (GC) per eye, for a total dose of 9 × 10 GC.

Three weeks post-injection, eyes were collected for episomal DNA extraction and PCR analysis to assess gene reconstitution. Junction-specific primers were designed to span the predicted synthetic introns formed by recombination between the 5′ and middle vectors (F1 and R1 primers) and between the middle and 3′ vectors (F2 and R2 primers) (black arrowheads in **Figure 1A**). PCR products with the expected sizes (624 bp for the 5’-middle junction and 510 bp for the middle-3’ junction) were detected exclusively in eyes that received the triple AAV-*EYS* vectors, regardless of whether the T2A-based (lanes 2-3) or IRES-based (lanes 4-5) 5’ vector was used (**Figure 3A**). No products were detected in uninjected control eyes (lane 1). The *EYS* expression plasmid (pcDNA3.1-hEYS) was used as a positive control (lane 6). The PCR products amplified from the control plasmid were slightly smaller (171 bp for the 5’-mid junction product and 166 bp for the mid-3’ junction product) than the corresponding amplicons from the AAV-injected eyes due to the lack of synthetic introns in the *EYS* expression plasmid. In contrast, PCR using primers F1 and R2 yielded no detectable amplicons under the same conditions (right-most panel in **Figure 3A**), confirming that no unintended recombination occurred directly between the 5’ and 3’ vectors.

**Figure 3.**
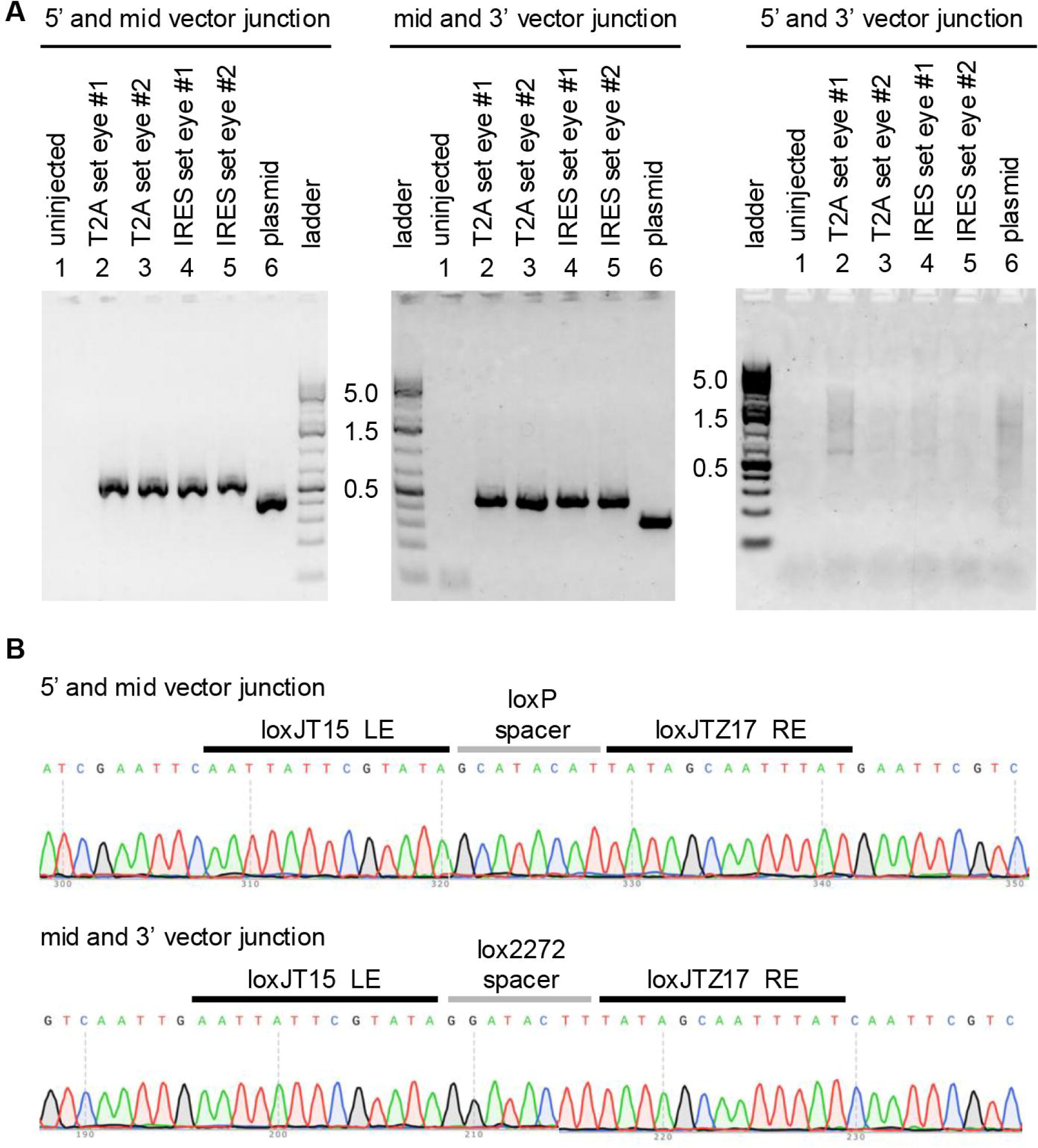
*In vivo* validation of *EYS* gene reconstitution in mouse retinas. **(A)** PCR analysis of junctional regions formed after vector recombination. Wild-type mice were injected subretinally with the complete AAV-*EYS* vector set at 1 month of age. Episomal DNAs were extracted 3 weeks post-injection from injected and uninjected eyes and used as templates for PCR amplification of the 5′-middle (with primers F1 and R1), middle-3′ (with primers F2 and R2), and 5’-3’ (with primers F1 and R2) junctions. The sizes of the 5′-middle and middle-3′ amplicons are 624 bp and 510 bp, respectively. No amplicons were detected Lane 1: uninjected eye; lanes 2-3: mice injected with the triple AAV-*EYS* set (with the T2A version of the 5′ vector); lanes 4-5: mice injected with the triple AAV-*EYS* set (with the IRES version of the 5′ vector); lane 6: positive control (full-length *EYS* plasmid). PCR products from the control plasmid are slightly smaller due to the absence of synthetic introns. **(B)** Sanger sequencing of PCR amplicons confirms correct junction formation. Sequences show the expected loxP (top) and lox2272 (bottom) spacers flanked by loxJT15 LE and loxJTZ17 RE, indicating precise recombination at both junctions.

To further validate recombination accuracy and amplicon identity, PCR products were Sanger-sequenced using the same forward primers employed for amplification (**Figure 3B**). As predicted, the 5′-middle junction contained the canonical loxP spacer flanked by the loxJT15 LE and loxJTZ17 RE elements (i.e., the lox15/17 double mutant), and the middle-3′ junction had the lox2272 spacer flanked by the same loxJT15 LE and loxJTZ17 RE elements (i.e., the lox15/17:2272 double mutant). Chromatograms showed no ambiguous peaks, indicating single-species amplification. These results confirm the fidelity of Cre-mediated recombination and verify that the orthogonal lox site pairs functioned as designed *in vivo*.

Protein expression was assessed by Western blotting of retinal lysates harvested 3 weeks post-injection. Consistent with the results from 293T cells, a ∼350 kDa EYS band was observed in all mouse eyes receiving the complete AAV-*EYS* vector set (**Figure 4**, black arrowhead in lanes 3-8). Both T2A-(lanes 3-5) and IRES-based (lanes 6-8) vectors facilitated *EYS* expression, indicating that either bicistronic configuration is sufficient for Cre expression and reconstitution. No EYS protein was detected in control uninjected retinas (lanes 1-2). A consistent non-specific band (open arrowhead) was observed across samples.

**Figure 4.**
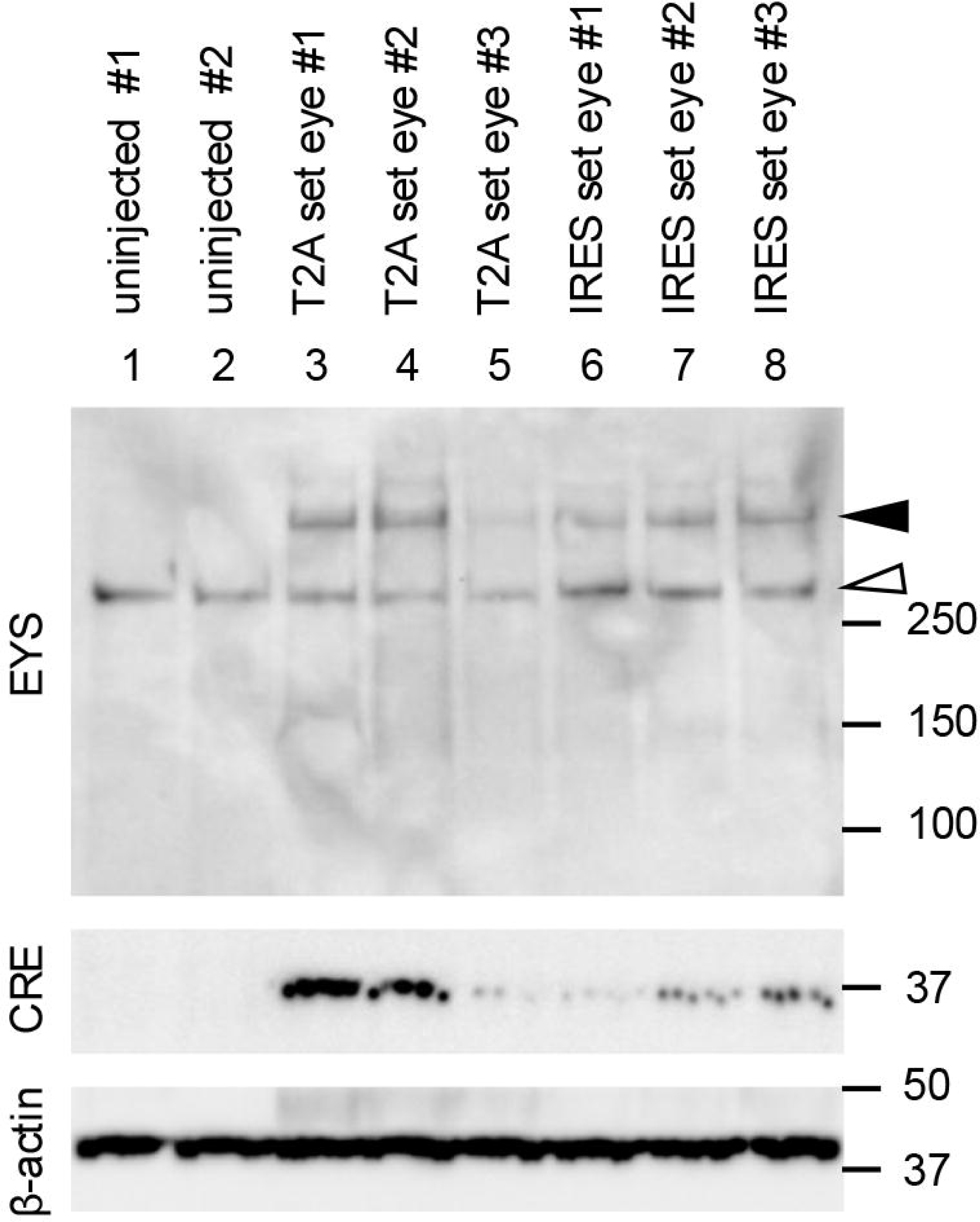
*In vivo* expression of full-length EYS protein in mouse retinas. Western blot analysis of retinal lysates collected 3 weeks post-injection. Lanes 1-2: uninjected controls; lanes 3-5: eyes injected with the complete tripartite AAV-*EYS* vector set containing the T2A-based 5′ vector; lanes 6-8: eyes injected with the complete tripartite AAV-*EYS* vector set containing the IRES-based 5′ vector. Full-length EYS is indicated by a black arrowhead; a non-specific band is marked with an open arrowhead. Each lane represents a different biological replicate.

Taken together, these results demonstrate that the tripartite vector system enables successful delivery, recombination, and expression of the full-length *EYS* gene in mouse photoreceptors, validating the feasibility of this approach *in vivo*.

## Discussion

The present study demonstrates a novel and effective strategy for delivering the full-length human *EYS* gene using a tripartite AAV system engineered with orthogonal lox sites and Cre recombinase. Given the large size of the *EYS* coding sequence (over 9 kb), traditional single-AAV vectors are incapable of accommodating this gene. Our approach overcomes this barrier by dividing the gene into three AAV-compatible fragments and reconstituting them *in situ* through unidirectional and site-specific recombination.

Compared to previous strategies such as dual/triple-AAV trans-splicing or split-intein systems,^22–24^ the Uni-STAR platform offers important advantages in efficiency, adaptability, and safety.

Whereas the AAV trans-splicing method relies on inefficient spontaneous recombination events, Cre recombinase catalyzes precise and efficient DNA recombination at pre-defined lox sites. Split-intein systems face constraints imposed by the positional requirements of intein motifs and by the stability and folding of truncated protein intermediates.^24,25^ Furthermore, because each vector in the split-intein approach continuously produces truncated proteins, there is a persistent risk of dominant-negative effects. By contrast, the Uni-STAR system restricts truncated protein production to the 5′ vector and only during the pre-recombination phase. As recombination progresses and the vectors are converted into full-length expression cassettes, truncated protein production ceases entirely. This feature provides an important safety advantage over the split-intein approach.

The lack of useful animal models for gene therapy preclinical studies poses a notable roadblock toward preclinical studies. In this regard, generation of EYS models in larger mammals such as rabbits, pigs, dogs, or non-human primates is urgently needed. The tripartite AAV system described here offers a powerful platform that can be directly applied to such models for preclinical validation.

A major challenge in the development of *EYS* gene therapies is the absence of suitable animal models for efficacy testing. While zebrafish mutants have yielded valuable insights into the developmental roles of *EYS*, they are not appropriate for evaluating mammalian gene therapy vectors. Rodents, including mice and rats, completely lack an *EYS* ortholog, making natural disease modeling impossible, and no mammalian models of *EYS*-associated retinopathies have been described to date. This lack of relevant models represents a critical roadblock to preclinical development. In this regard, generation of *EYS* disease models in larger mammals—such as rabbits, pigs, dogs, or non-human primates— is urgently needed. The tripartite AAV system described here offers a ready-to-use platform that can be applied directly to such models, enabling rigorous preclinical validation and accelerating the path toward clinical translation.

In conclusion, this study demonstrates that the Uni-STAR-based tripartite AAV system provides an efficient and versatile strategy for delivering large genes such as *EYS*. By utilizing unidirectional, site-specific recombination, this approach overcomes the principal obstacle to *EYS* gene therapy and establishes a broadly applicable framework for treating not only inherited retinal diseases but also diverse genetic disorders caused by genes that exceed the natural packaging capacity of AAV.

## Conflict of Interest Statement

S.S. and P.D. are co-inventors on a patent application (19/124,578, filed by the University of Iowa) titled “Method to deliver large genes using virus and a DNA recombination system.” The remaining authors declare no competing interests.

## Authorship Confirmation

Conception and study desing: S.S., Plasmid construction: K-D.R., P.D., Mouse colony maintenance: C.B., Investigation: K-D.R., P.D., C.B., S.S., Validation: K-D.R., P.D., Data curation: S.S., Writing – Original Draft: S.S., Writing – Review & Editing: K-D.R., P.D., Visualization: S.S., Supervision: S.S., Project administration: S.S., Funding acquisition: S.S.

## Funding Statement

This work was supported by National Institutes of Health grant R01-EY034176 (S.S.), Retina Research Foundation Pilot grant (S.S.), Flagellar Vision Foundation (S.S.).

